# Dissociation of SYNGAP1 Enzymatic and Structural Roles: Intrinsic Excitability and Seizure Susceptibility

**DOI:** 10.1101/2025.01.14.633019

**Authors:** Julia Brill, Blaise Clarke, Ingie Hong, Richard L. Huganir

## Abstract

SYNGAP1 is a key Ras-GAP protein enriched at excitatory synapses, with mutations causing intellectual disability and epilepsy in humans. Recent studies have revealed that in addition to its role as a negative regulator of G-protein signaling through its GAP enzymatic activity, SYNGAP1 plays an important structural role through its interaction with post-synaptic density proteins. Here, we reveal that intrinsic excitability deficits and seizure phenotypes in heterozygous Syngap1 knockout (KO) mice are differentially dependent on Syngap1 GAP activity. Cortical excitatory neurons in heterozygous KO mice displayed reduced intrinsic excitability, including lower input resistance, and increased rheobase, a phenotype recapitulated in GAP-deficient Syngap1 mutants. However, seizure severity and susceptibility to pentylenetetrazol (PTZ)-induced seizures were significantly elevated in heterozygous KO mice but unaffected in GAP-deficient mutants, implicating the structural rather than enzymatic role of Syngap1 in seizure regulation. These findings highlight the complex interplay between SYNGAP1 structural and catalytic functions in neuronal physiology and disease.

**Significance Statement:** Mutations in the *SYNGAP1* gene are a major cause of intellectual disability, autism, and epilepsy. The SYNGAP1 protein is an important constituent of postsynaptic specializations, and two distinct functions have been characterized: a structural function, carried by its C-terminal PDZ domain, that organizes the composition of the postsynaptic density, and an enzymatic function, in which its GAP domain negatively regulates small GTPases. So far, no electrophysiological/behavioral phenotype of SYNGAP1 has been directly linked to the GAP catalytic activity. Here, we describe that while the GAP catalytic activity does not contribute to the increased seizure susceptibility seen in SYNGAP1 haploinsufficiency, it does regulate the intrinsic excitability of upper lamina pyramidal cells.

## Introduction

SYNGAP1 is a synaptic GTPase-activating protein (GAP) critical for synaptic plasticity, learning, memory, and cognition. SYNGAP1 modulates Ras and Rap signaling via its GTPase-activating (GAP) activity, but is also one of the most abundant proteins at excitatory synapses and may be important in the dynamic structural integrity of the postsynaptic density (PSD) (1-5).

Studies using knockout (KO) mice have demonstrated that a heterozygous knockout of Syngap1 leads to significant impairments in long-term potentiation (LTP), premature dendritic spine maturation, and learning deficits (6-8). Subsequent research revealed spontaneous seizure activity and heightened seizure susceptibility in these mice (9,10). Homozygous knockout mice, on the other hand, do not survive beyond the first few days after birth (6-8), a period coinciding with a marked increase in Syngap1 expression in the brain (11, 12). Reduced intrinsic excitability in cortical excitatory neurons was found in heterozygous Syngap1 knockout mice, a phenotype associated with higher rheobase, hyperpolarized resting membrane potential (RMP), and decreased input resistance (13). Importantly, mutations in the *SYNGAP1* gene in humans result in a spectrum of neurodevelopmental disorders, including severe intellectual disability, autism spectrum disorders, and epilepsy (14-16). These neuronal and systemic abnormalities in mice and humans highlight Syngap1’s essential role in postnatal brain development and synaptic regulation, underscoring its importance as a key molecular hub in synaptic function.

Recent studies suggest that SYNGAP1 may independently regulate synaptic strength and plasticity through structural mechanisms. At excitatory synapses, SYNGAP1 undergoes liquid-liquid phase separation (LLPS) with PSD-95 (11, 17,18), a process that can exclude other proteins from liquid-like concentrates with interacting partners. The high abundance of SYNGAP1 and its occupation of the PDZ domain of PSD-95 prevent binding and synaptic incorporation of AMPA receptor/TARP complexes (11, 19, 20), providing a brake on synaptic strengthening. The subsequent dispersion of SYNGAP1 from synapses during plasticity (21) can thus relieve this steric blockade and allow other molecules to take up the synaptic ‘slot’, leading to synaptic strengthening. Importantly, mice with mutations inactivating the GAP enzymatic function of SYNGAP1 did not show synaptic plasticity impairments, behavioral impairments, or postnatal mortality, suggesting that its structural properties may underlie many of SYNGAP1’s function in neurons (22). However, the relative contributions of its GAP activity and structural function to seizure pathophysiology and intrinsic excitability remain unclear.

This study investigates whether these deficits are mediated by SYNGAP1 GAP activity. We employed GAP-deficient Syngap1 mutant mice to delineate the enzymatic and structural contributions of Syngap1 in modulating intrinsic neuronal excitability and seizure susceptibility. Our findings demonstrate that GAP activity is essential for maintaining intrinsic excitability, whereas seizure regulation is primarily driven by the structural role of Syngap1.

## Results

### Intrinsic Excitability is Regulated by the Enzymatic Function of Syngap1

Layer 2/3 pyramidal cells in somatosensory cortex of Syngap1 heterozygous knockout mice have reduced intrinsic excitability (13). Since Syngap1 knockout eliminates both its structural and enzymatic function, it remained unclear which of these factors underlies the reduction in intrinsic excitability. To clarify, we measured the intrinsic excitability of pyramidal cells in somatosensory cortex of mice with targeted point mutations abolishing Syngap1’s Ras-GTPase activity while leaving its structural function unaffected (GAP* mice) (22).

Intrinsic excitability of pyramidal cells undergoes profound developmental changes (23-25); therefore, we made sure to always use age-matched mice and keep close track of developmental changes observed in wildtype (WT, +/+) mice. The developmental trajectory of intrinsic excitability components measured in this study in wildtype mice can be found in (*SI Appendix*, Fig. S1).

We found that the intrinsic excitability of layer 2/3 pyramidal cells was reduced in the GAP*/GAP* mice, similar to the knockout mice. In slices from GAP*/GAP* mice aged P38-48, layer 2/3 pyramidal cells had significantly reduced spike frequency in response to depolarizing current injections (FIG. 1*A, C*, and significantly increased “rheobase”, i.e., the lowest current injection that elicited spiking (Table 1). Cells in slices from +/GAP* mice displayed intermediate properties but did not differ statistically from cells of +/+ or GAP*/GAP* littermates. These effects mirror results obtained from age-matched heterozygous knockout mice (Fig 1*D*; *SI Appendix*, Fig. S2H). There, pyramidal cells from mutant mice likewise had a 25-30% increased rheobase (Table 1) and reduced spike frequency, similar to the results reported by Michaelson et al (13). Cells from +/*, */* and +/− animals also had a more depolarized spike threshold compared to littermate controls (Fig. 1*B*) which might directly contribute to their increased rheobase (Table 1). To further characterize the developmental trajectory in GAP* mice, we also measured intrinsic excitability in slices from younger animals, aged P11-14. Intrinsic excitability in layer 2/3 pyramidal cells decreases during development (*SI Appendix*, Fig. S1), and accordingly, we observed a higher spike frequency and lower rheobase in the younger compared to older wildtype mice. Just like in the older age group, pyramidal cells of younger GAP* mice displayed reduced excitability compared to age-matched littermates, with a near doubling of rheobase and decreased firing frequency in GAP*/GAP* mice and intermediate values in +/GAP* (Fig. 1*E, G*; Table 1).

**Figure 1.**
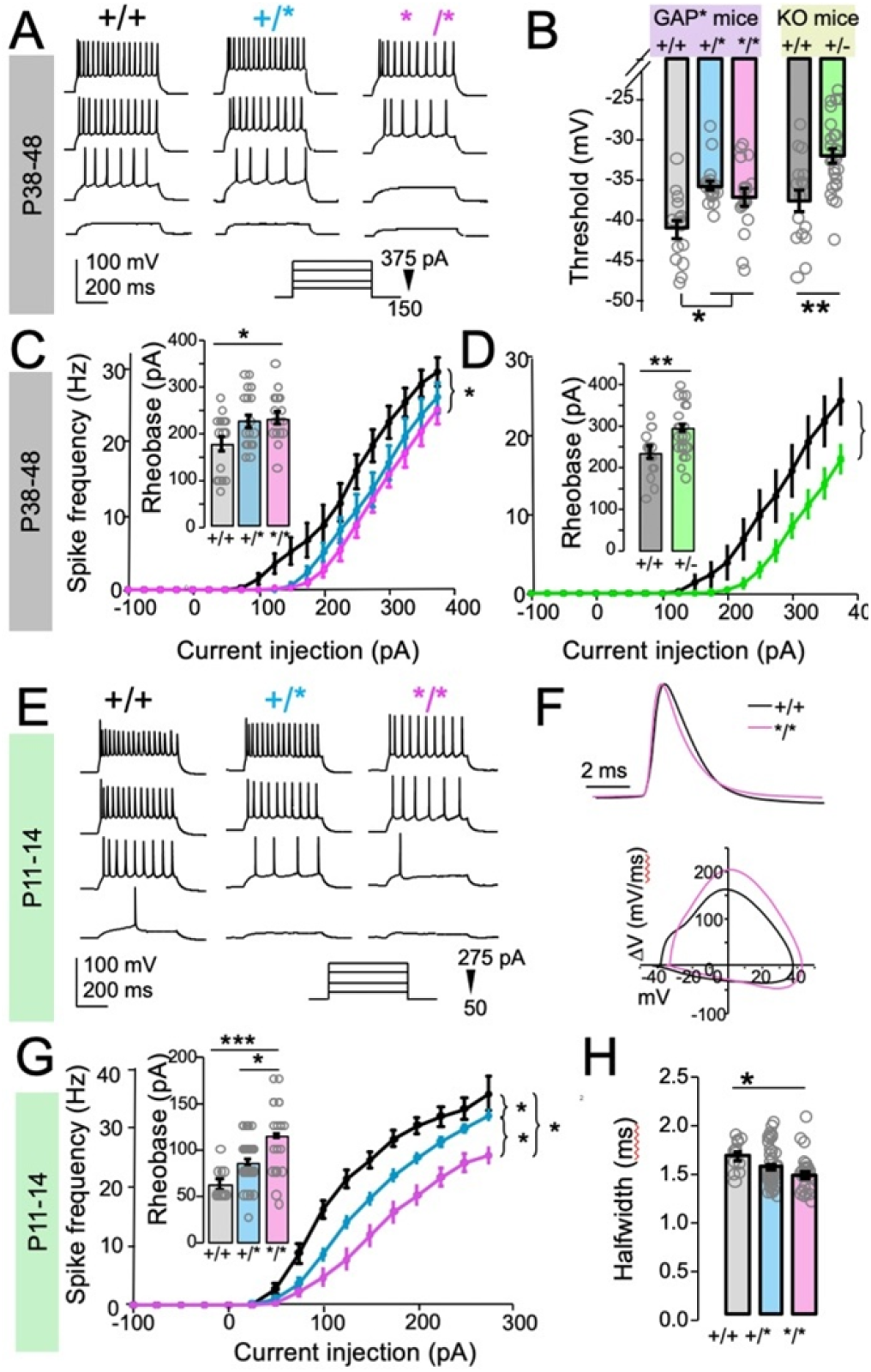
Layer 2/3 pyramidal cells in Syngap1 +/GAP* and GAP*/GAP* mice have reduced intrinsic excitability, similar to heterozygous KO mice. (A-D) Data from P38-48 GAP* and KO mice. (A) Example current clamp traces in from 500 ms current injections in +/+ (WT), */GAP* (+/*) and GAP*/GAP* (*/*) cells, showing reduced spiking and increased rheobase in +/* and */* cells. Current injections of +150, +225, +300 and +375 pA (bottom to top). (B) Summary graphs showing spike threshold in pyramidal cells from +/+, +/* and */* mice, as well as +/+ and +/− mice. (C) Input/output curves showing spike frequency in response to hyperpolarizing and depolarizing current injections for GAP* mice. Inset: Rheobase in +/+,+/* and */* cells. (D) Input/output curves showing spike frequency in response to hyperpolarizing and depolarizing current injections for KO mice. Inset: Rheobase in +/+ and +/−cells. (E-H) Data from P11-14 GAP* mice. (E) Example current clamp traces in from 500 ms current injections in +/+ (WT), */GAP* (+/*) and GAP*/GAP* (*/*) cells, showing reduced spiking and increased rheobase in +/* and */* cells. Current injections of +50, +125, +200 and +275 pA (bottom to top). (F) Top: Representative action potentials from WT (black) and */* (magenta) mice, scaled to peak and aligned to threshold. Bottom: Phase plot (V vs ΔV) from the action potentials shown in the left. The action potential from the */* cell has a shorter halfwidth and both faster rise and decay phases. (G) Input/output curves showing spike frequency in response to hyperpolarizing and depolarizing current injections for GAP* mice. Inset: Rheobase in +/+,+/* and */* cells. (H) Summary graphs showing spike halfwidth in pyramidal cells from +/+, +/* and */* mice. *, **, *** indicate p<0.05, p<0.01, p<0.005, respectively. Unpaired t-test for comparing +/+ vs. +/−; ANOVA for comparing +/+ vs +/* vs */*.

**Table 1.**
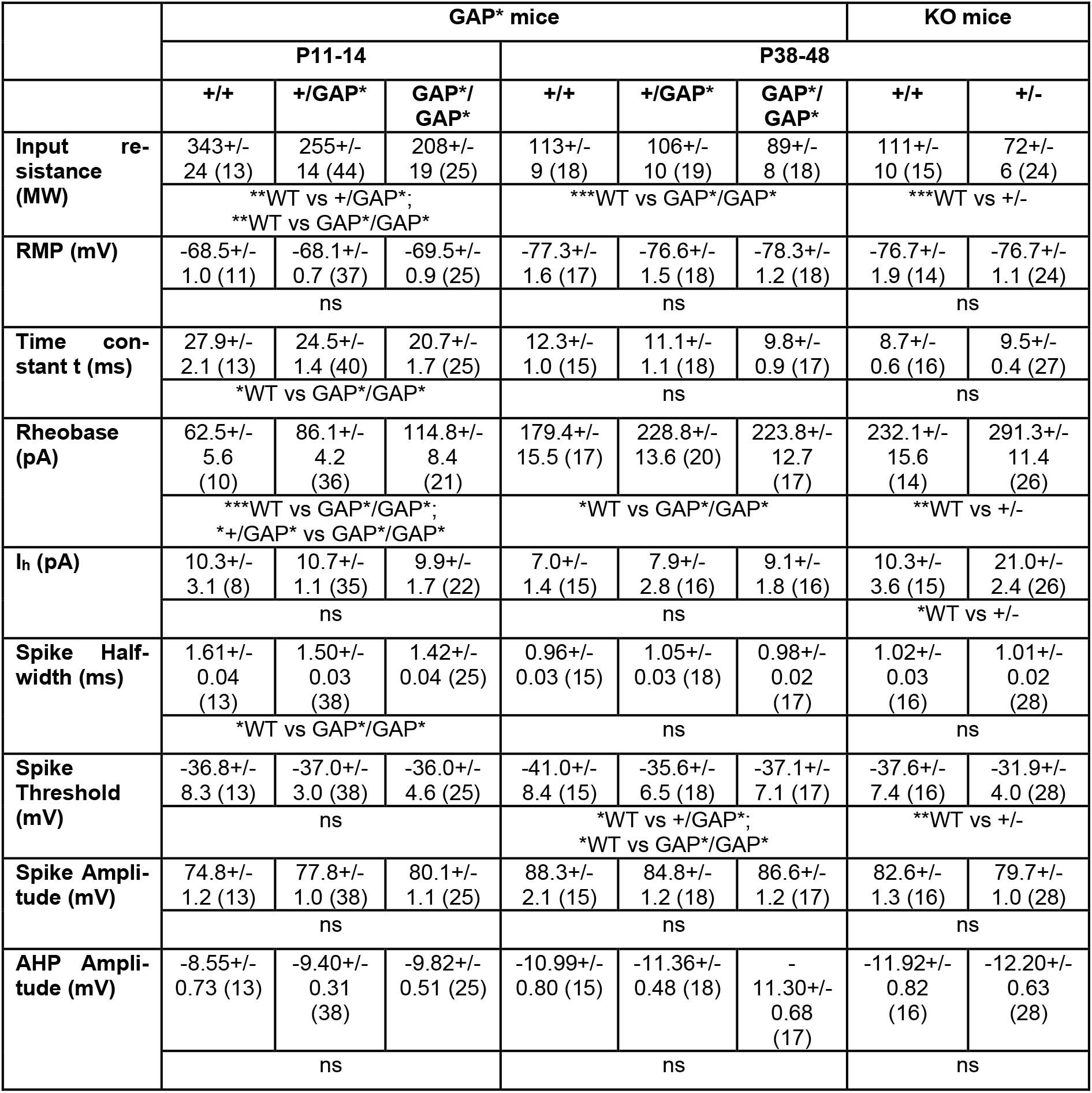
Summary of the intrinsic properties measured in this study. Data is given as mean +/− SEM (sample size). *, **, *** indicate p<0.05, p<0.01, p<0.005, respectively. Unpaired t-test for comparing +/+ vs. +/−; ANOVA for comparing +/+ vs +/* vs */*.

### Increased Potassium Conductance is Associated with Loss-of-Function of Syngap1

To further examine action potential properties and kinetics, we chose the first spike evoked in each cell during the administration of depolarizing current injections. There were no significant differences in spike amplitude and afterhyperpolarization in cells from either age group or strain (*SI Appendix*, Fig. S2). Spike halfwidth was reduced in cells from younger GAP*/GAP* mice, but not in either genotype in the older age group (Fig. 1*F, H*; *SI Appendix*, Fig. S2).

We also measured passive membrane properties, i.e. resting membrane potential (RMP), input resistance, and membrane time constant, in these cells. There were no significant genotype differences in RMP in either age group (Fig. 2*A*). Input resistance was measured in voltage clamp mode while holding cells at −60 mV and delivering brief hyperpolarizing voltage steps (to −65 mV). We found significant reductions in input resistance in both age groups and both mutant strains (Fig. 2*B*). In P11-14 cells from GAP* mice, input resistance was reduced in heterozygous as well as homozygous animals (Table 1), while in the older age group, only the reduction in homozygous mutants reached statistical significance, with heterozygous mice showing intermediate values. Input resistance was also reduced in cells from +/− mice. The membrane time constant was significantly shorter in cells from younger GAP*/GAP* mice but did not differ from wildtype in the older GAP* or knockout mice (Fig. 2*C*). The only difference between the two strains that reached statistical significance was an somewhat increased I_h_ in +/−, but not GAP* mice (Table 1).

**Figure 2.**
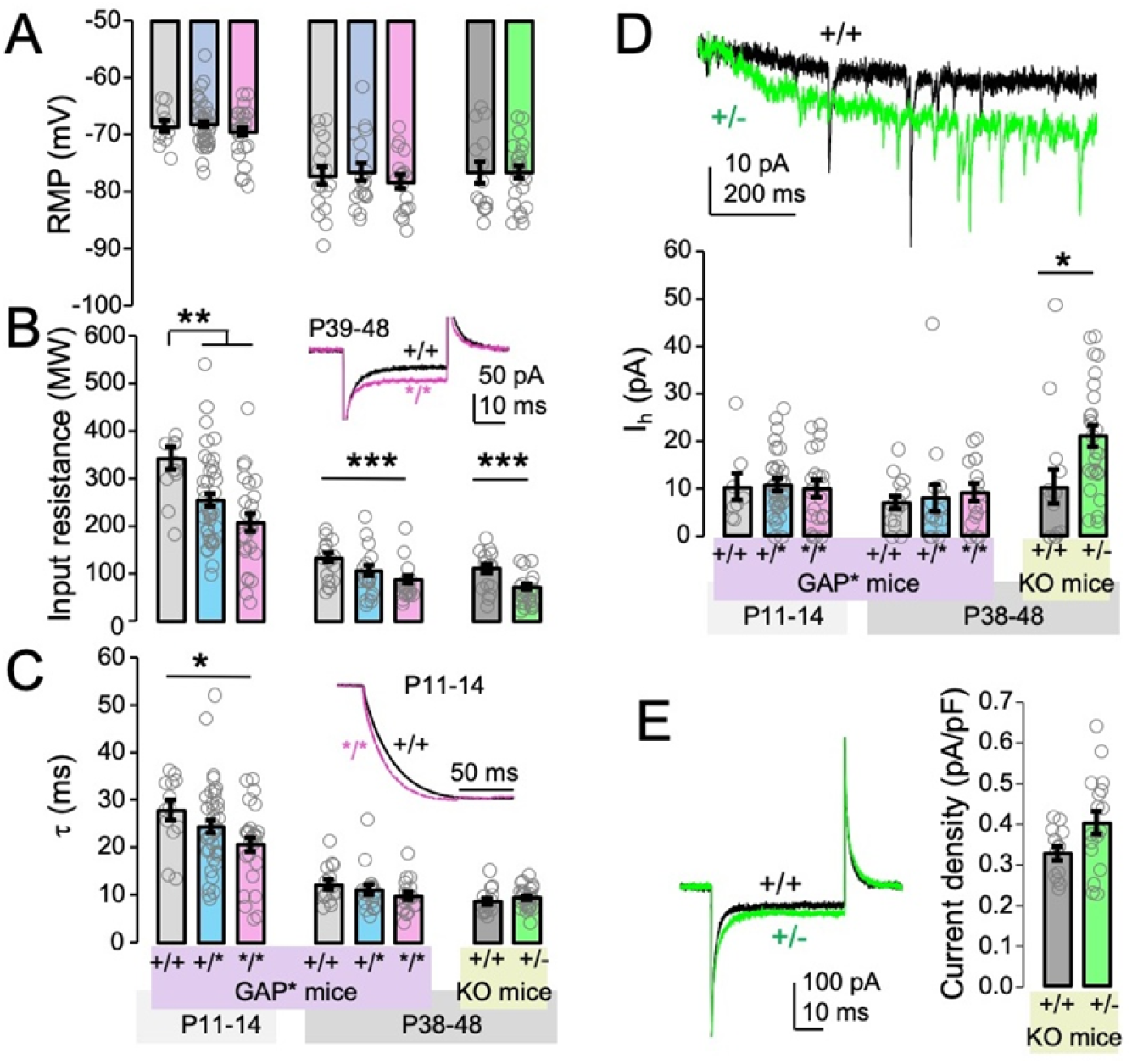
Decreased input resistance is the likely cause of reduced excitability. (A-C) Passive membrane properties. Summary graphs showing RMP (A), input resistance (B) and membrane time constant (C) in pyramidal cells from younger and older wt, +/* and */* mice, as well as wt and +/− mice. Insets in B and C show representative traces from wt and */* mice highlighting decreased input resistance in */* (B) and shorter membrane time constant in */* cells the younger age group (C). (D) I_h_. Top: Exemplar traces from P38-48 WT (black) and +/− (green) mice showing currents in response to a hyperpolarizing voltage step (−60 to −85 mV) to measure I_h_. A larger current is observed in the +/− cell. Bottom: Summary graphs showing I_h_ in pyramidal cells from younger and older wt, +/* and */* mice, as well as wt and +/− mice. The only significant difference is observed in cells from +/− mice. (E) Leak potassium currents. Left: Example response to −60 to −65 mV voltage step in the presence of TTX and CdCl_2_ in cells from wt (black) and +/− (green) animals, aged P30-35. Larger current is evoked in the +/− cell, indicating higher leak currents. Cell capacitance was measured as the area under the curve of the second capacitive current transient (which is similar in both cells). Right: Summary graphs showing leak current density (current/capacitance) in wt and +/− cells. Current density is increased in +/− cells, consistent with larger leak potassium currents. *, **, *** indicate p<0.05, p<0.01, p<0.005, respectively. Unpaired t-test for comparing +/+ vs. +/−; ANOVA for comparing +/+ vs +/* vs */*.

Lower input resistance alongside decreased excitability could be due to increased leak potassium conductance (26). Therefore, we measured the leak potassium current density using voltage steps from −60 to −65 mV (as was done to measure input resistance) in cells from +/− mice the presence of TTX and CdCl_2_ to block sodium and calcium channels, respectively. Cells from +/− mice had a significantly higher current density than cells from wildtype littermates (+/+: 0.326+/−0.017 pA/pF, n=13; +/−: 0.401+/− 0.029 pA/pF, n=17; p<0.05), consistent with increased leak potassium currents (Fig. 2*E*).

Together, our findings show that layer 2/3 pyramidal cells in both GAP* and Syngap1 knockout mouse strains display decreased input resistance, suggesting this is likely an underlying cause of their reduced intrinsic excitability.

### PTZ-Induced Seizures in Syngap1 Mutants Reveal Differential Effects of Structural and Enzymatic Functions

We assessed the severity and susceptibility of pentylenetetrazol (PTZ)-induced seizures in Syngap1 heterozygous knockout (KO) and GAP domain-mutant (GAP*) mice to discern the contributions of SYNGAP1’s structural and enzymatic functions to seizure phenotypes. Seizure severity was quantified using a modified Racine scale scored during sequential PTZ injections (27), while seizure susceptibility was evaluated as the cumulative percentage of mice remaining free of tonic-clonic seizures at each dose.

In Syngap1 KO (+/−; n=10) mice, the maximum Racine scale scores were significantly elevated compared to wildtype littermate (+/+; n=14) controls, typically reaching the tonic-clonic seizure threshold (score 6) after the first PTZ dose (Fig. 3*A*). The survival analysis further demonstrated that Syngap1 KO mice exhibited a marked reduction in resistance to tonic-clonic seizures, with the majority succumbing to seizures earlier than WT mice (Fig. 3*B*). These findings confirm heightened seizure susceptibility and severity in Syngap1 haploinsufficient mice.

**Figure 3.**
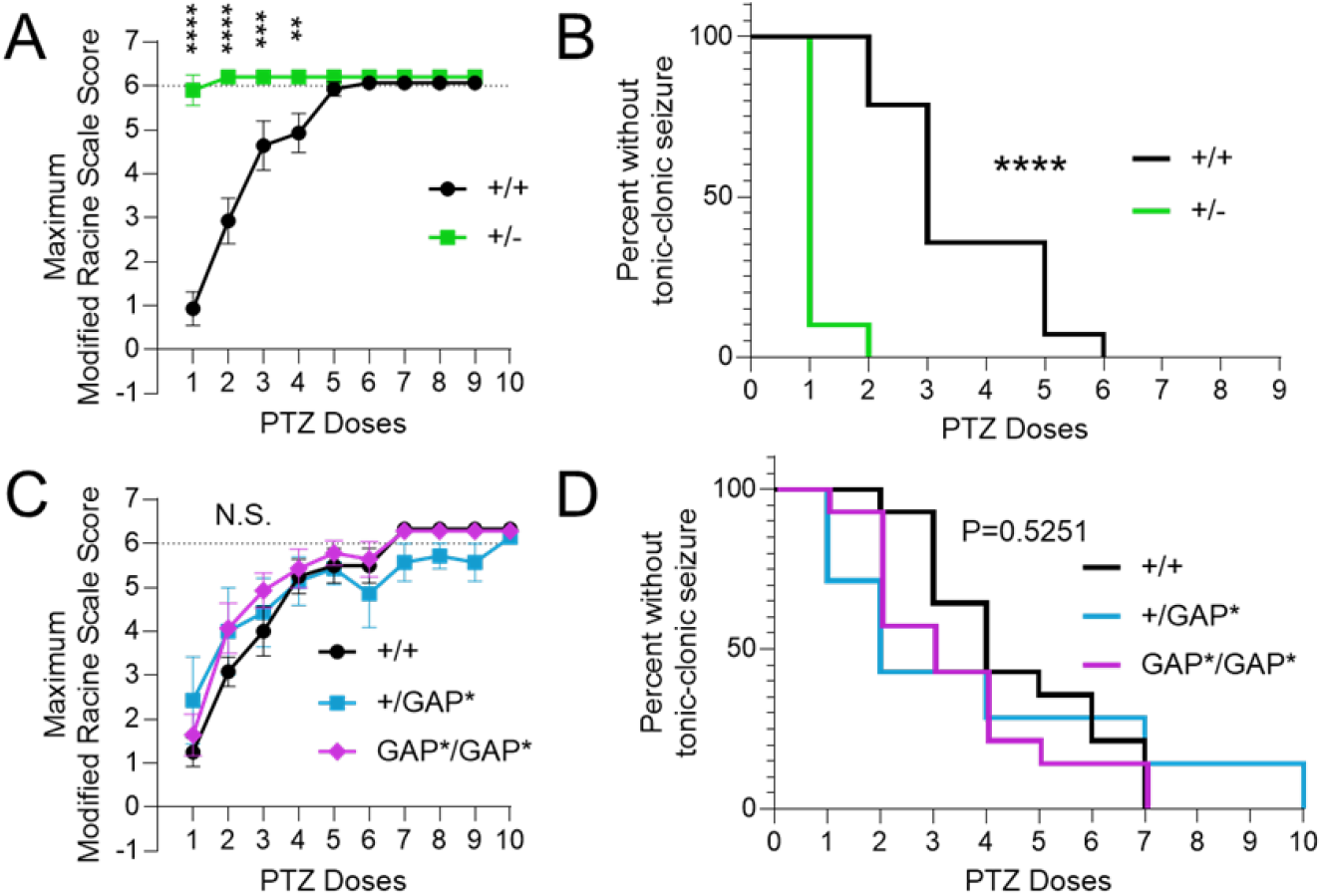
PTZ-induced seizure severity and susceptibility are heightened in Syngap1 heterozygous knockout but not +/GAP* and GAP*/GAP* mice. (A,C) Modified Racine scale scores were used to evaluate PTZ-induced seizures. Maximum scores following sequential PTZ administration (40 mg/kg initial dose, 10 mg/kg every 5 minutes, until a score of 6 or higher was observed) in Syngap1 KO (A) and GAP* (C) mutants were plotted. Dotted line indicates tonic-clonic seizure threshold. Data points show mean ± SEM. *, **, ***, **** indicate p<0.05, p<0.01, p<0.001, p<0.0001, respectively. Two-way ANOVA (genotype x dose) with Šídák’s multiple comparisons test was used to compare seizure severity for each PTZ dose. (B,D) Survival plots showing percentage of mice without tonic-clonic seizures for Syngap1 KO (B) and GAP* (D) mutants after each administration. The Log-rank (Mantel-Cox) test was used to compare survival curves.

In contrast, GAP domain mutants (+/GAP*, n=6; GAP*/GAP*, n=14) did not exhibit such heightened seizure susceptibility or severity. The maximum Racine scale scores of these mice were comparable to WT littermates (n=12), and survival analyses indicated no significant difference in resistance to tonic-clonic seizures across genotypes (Figs. 3*C*, 3*D*). These results suggest that the increased seizure susceptibility observed in Syngap1 KO mice is not dependent on SYNGAP1 GAP activity but rather on its structural functions.

Together, these data underscore a dissociation between the structural and enzymatic roles of SYNGAP1 in regulating seizure phenotypes. While the structural contributions are critical for maintaining normal seizure thresholds, GAP activity appears dispensable for this function.

## Discussion

SYNGAP1 is one of the most abundant postsynaptic proteins in excitatory synapses of glutamatergic neurons (28). So far, two distinct functions have been identified: a *structural role* via its PDZ domain, controlling which proteins enter and leave the postsynaptic density (19, 21), and an enzymatic activity, i.e. a GTPase activating domain (GAP) that negatively regulates several small GTPases including Ras, Rap and Rab (4, 29-31).

We have developed a mouse strain with two amino acid substitutions within the GAP domain that abolish its enzymatic activity (7, 32, 33) without affecting protein expression and targeting (GAP* mice) (22). The characterization of Syngap1 GAP mutant mice so far has revealed that homozygotes are viable and fertile - unlike homozygous Syngap1 KO mice which die perinatally (8). Although GAP*/GAP* mice show minor differences in dendritic spine size, unlike Syngap1 +/− mice, they do not have overt behavioral deficits and impaired LTP (22).

In this study we extend our characterization of GAP* mice by examining cellular and network excitability. Previous studies have shown that Syngap1 haploinsufficient mice have an increased seizure susceptibility (9, 34) and altered excitability in several neuron types, including upper lamina cortical pyramidal cells (13), and striatal projection neurons (35). We found that GAP* mutant mice, unlike +/− mice, did not exhibit increased susceptibility to PTZ-induced seizures. However, both Syngap1 mutant strains had the same deficit in intrinsic excitability of layer 2/3 pyramidal cells in somatosensory cortex. This is the first electrophysiological phenotype that is attributable to the GAP catalytic activity of Syngap1. The only other mutation in rodents that targets the GAP domain is a larger deletion in rats, which encompasses most of the GAP and preceding C2 domains (36). Unlike GAP* mice, these rats display a phenotype similar to that of haploinsufficient mice, suggesting that this larger deletion might affect protein expression, localization, and/or structural function.

We were able to show that the intrinsic excitability of upper lamina pyramidal cells in somatosensory cortex of GAP* mutant mice was reduced to a similar extent as cells from Syngap1 haploinsufficient mice. Cells from both strains had an increased rheobase, reduced overall spike output in response to depolarizing current injections, and reduced input resistance. The reduction in input resistance sufficiently explains the reduced firing and increased rheobase of those cells. In all of these points, GAP* and knockout mutant cells were alike. Thus, we have positively identified an electrophysiological function of SYNGAP1 that is due to its enzymatic rather than structural activity.

Some evidence suggests that increased leak potassium currents could be responsible for the decreased input resistance and excitability. Leak potassium channels are major contributors to neuronal input resistance and are master regulators of neuronal excitability and have a large influence over neuronal input resistance, stabilize the resting membrane potential and are, among other factors, modulated by small GTPases (26, 37-39). They are therefore a plausible underlying cause of the observed phenotype.

One difference between the two strains with regards to intrinsic excitability was increased hyperpolarization induced cation current (I_h_) in Syngap1 knockouts, but not in GAP* mutants. I_h_, carried by HCN channels, tends to promote spiking via rebound excitation (40), and is largely inactive above –60 mV. Thus, it is unlikely to be the cause of the decreased intrinsic excitability of +/− cells. In light of the increased seizure susceptibility of haploinsufficient mice, it is interesting to note that HCN channel inhibition can suppress absence seizures (41).

It seems paradoxical that reduced neuronal excitability would be found in animals with increased seizure susceptibility. The dissociation of excitability and seizure phenotypes in our results show that the reduced excitability is not causal to the increased seizure susceptibility, which would have been difficult to rationalize. Rather, the seizure phenotype in haploinsufficient mice exists despite, not because of, the reduced neuronal excitability. It should be noted though that we studied only cell type, layer 2/3 pyramidal cells, and that other cell types can be affected differently. For example, striatal indirect pathway neurons (iSPNs) have increased intrinsic excitability in Syngap1^+/−^ mice (35). Additionally, seizures are a network rather than cell intrinsic phenomenon and reduced intrinsic excitability of glutamatergic neurons could conceivably lead to less activation of GABAergic neurons and thus disinhibition under certain circumstances. Importantly, five human *SYNGAP1* single nucleotide variant carriers with GAP-disabling mutations were found not associated with any neurological or mental diagnosis (22), suggesting that the loss of GAP function does not lead to seizures in humans. This is consistent with our results in mice and indicates that future therapeutic strategies for the treatment of SYNGAP1 haploinsufficiency should include means to rescue the structural function of SYNGAP1.

## Acknowledgements

We would like to thank Yoichi Araki, Richard C. Johnson, Austin Graves, Sarah Rodriguez, and Ashley Irving for technical support, Qianwen Zhu for helpful discussions and David Linden for his unwavering support. Funding: This work was supported by National Institutes of Health grants R37NS036715 (RLH) and U01DA056556 (RLH and IH).

## Author contributions

Conceptualization: JB, IH, and RLH; Methodology: JB, IH, BC, and RLH; Investigation: JB, IH, BC and RLH; Validation: JB, IH and RLH; Visualization: JB, BC, and IH; Formal analysis: JB, IH, and BC; Data curation: JB and IH; Resources: IH and RLH; Supervision: IH and RLH; Project administration: IH and RLH; Funding acquisition: IH and RLH; Writing: JB, IH, BC and RLH.

## Materials and Methods

All the experiments reported in this study adhered to the ethical regulations and guidelines of the National Institute of Health Guide for Care and Use of Laboratory Animals. The experiments involving mice were approved by the Institutional Animal Care and Use Committee at Johns Hopkins University School of Medicine under the study protocol numbers: M023M70, MO22M278, and MO23M52.

### Mice

All animal experiments adhered to the guidelines of the National Institute of Health and were approved by the Institutional Animal Care and Use Committee at Johns Hopkins University School of Medicine. Mice were maintained on a 14h light/10h dark cycle. Both male and female mice were used, and no obvious differences between the sexes were noted. All mice were group-housed in pathogen-free facilities with regulated temperature and humidity and given *ad libitum* access to food and water. All mice used were seemingly free of infection, health abnormalities or immune system deficiencies and were employed independently of their gender. None of the mice used had been used for previous experiments. The date of the vaginal plug detection was designated E0.5, and the date of birth P0. *Syngap1*^*+/−*^ heterozygous knockout mice (8), *Syngap1* GAP mutants (*+/GAP*, GAP*/GAP\**) (22), and WT (*Control*) littermates were maintained on a mixed background of C57/B6J and 129/SvEv background.

### Seizure Severity and Susceptibility Measurements

Seizure susceptibility was assessed by administering pentylenetetrazol (PTZ) to mice. PTZ was diluted to 2 mg/mL in 0.9% sterile saline and given to mice via intraperitoneal (IP) injection. Before PTZ administration, mice were habituated in a recording chamber for five minutes. An initial dose of 40 mg/kg PTZ was then given, followed by additional doses of 10 mg/kg every five minutes until a tonic-clonic seizure was observed. Researchers were blinded to the genotype of the animals. Severity of seizure was quantified using a modified Racine scale (34) ranging from −1 (baseline), 0 (whisker trembling), 1 (sudden behavioral arrest), 2 (facial jerks), 3 (neck jerks), 4 (sitting: clonic seizures), 5 (lying on belly), 6 (lying on side or wild jumping) to 7 (wild jumping, respiratory arrest, death), along with video monitoring of the mice’s behavior. A score of 6 is defined as a tonic-clonic seizure, the goal score for this experiment. If a mouse progressed to a score of 6 or higher during an administration, the previous dose was recorded as the final one.

Following the experiment, mice that did not undergo respiratory arrest were euthanized via isoflurane inhalation. Brain tissue was then collected and preserved by flash-freezing in dry ice, then storing at −80C for later experiments. Video recordings of the assessments were captured and stored for additional post-hoc analysis.

### Electrophysiology

#### Slice preparation

To prepare brain slices, mice were deeply anesthetized using 5% isoflurane, brains were removed and submerged in ice-cold NMDG-HEPES-aCSF containing, in mM: 92 NMDG, 2.5 KCl, 1.25 NaH_2_PO_4_, 30 NaHCO_2_, 20 HEPES, 25 glucose, 0.5 CaCl_2_, 10 MgSO_4_; titrated to pH 7.35 with HCl and bubbled with carbogen (95% O_2_/ 5% CO_2_). Coronal neocortical slices (350 μm) were sectioned on a VT 1200S Vibratome (Leica) at 4°C in ice-cold NMDG-HEPES-aCSF solution, and then transferred into a recovery chamber filled with 150 ml of recovery aCSF containing, in mM: 126 NaCl, 2.5 KCl, 1.25 NaH_2_PO_4_, 26 NaHCO_3_, 2 MgCl_2_, 2 CaCl_2_, 10 glucose, 2 thiourea, 5 Na-ascorbate, 3 Na-pyruvate; warmed to 32°C and bubbled with 95% O_2_:5% CO_2_. After 30 minutes of recovery, slices were incubated at room temperature for a minimum of 45 minutes before the start of experiments.

#### Whole cell recordings

Slices were transferred into a submerged recording chamber and continually superfused at a rate of ∼2ml/min with aCSF containing, in mM: 126 NaCl, 2.5 KCl, 1.25 NaH_2_PO_4_, 26 NaHCO_3_, 2 MgCl_2_, 2 CaCl_2_, and 10 glucose, equilibrated with 95% O_2_:5% CO_2_. All experiments were conducted at 32°C. Signals were acquired using an Axon Multiclamp 700B amplifier, Digidata 1440 A digitizer and pClamp software, sampled at 10 to 100 kHz, and filtered at 3 to 30 kHz. Whole-cell voltage-clamp and current-clamp recordings were obtained using borosilicate glass electrodes with a tip resistance of 2–4 MΩ. The pipette solution for recording potassium currents contained, in mM: 140 KCl, 5 EGTA, 10 HEPES, 2.5 MgCl_2_. The pipette solution for all other experiments contained, in mm: 120 K-gluconate, 11 KCl, 2 MgCl_2_, 2 CaCl_2_, 10 HEPES, 10 EGTA, pH 7.3, adjusted with KOH. Neuronal firing frequency were assessed in current-clamp mode by giving a series of hyperpolarizing and depolarizing current injections (0.5 s each), typically −100 to +300 pA in 25 pA increments. No holding current was applied. Action potential waveforms were quantified for the first spike evoked at the lowest current that resulted in spiking. Resting membrane potential was determined from a 5-30 s gap-free current clamp recording. Input resistance and capacitance were determined in voltage clamp mode by applying a 50 ms voltage step from −60 to −65 mV. 30 sweeps recorded at 30 kHz were averaged. The membrane time constant t was determined by fitting a first order exponential decay function (Clampfit) to the voltage recorded in response to a 100 pA hyperpolarizing current injection. To determine I_h_, a 500 ms step from −60 to −85 mV was applied in voltage-clamp mode and the difference in current at the beginning and end of the step was measured. Leak potassium currents were measured in voltage clamp mode using the same protocol as for measuring input resistance, but in the presence of 1 mM TTX to block sodium currents and 100 mM CdCl_2_ to block calcium currents. Data were analyzed in Clampfit.

### Statistical analysis

Statistical analyses of the electrophysiology and seizure behavior were done using GraphPad Prism and Microsoft Excel software. Statistically significant differences were assessed by Student’s unpaired t-test two compare two groups or ANOVA to compare three groups. Two-way ANOVA (genotype x dose) with Šídák’s multiple comparisons tests was used to compare seizure severity for each PTZ dose. Log-rank (Mantel-Cox) test was used to compare survival curves. P<0.05 was considered a significant difference. All values represent individual animals mean ± standard error of the mean (SEM). The statistical test used, and the statistical significance are indicated in figure legends.

## Supporting Information for

Dissociation of SYNGAP1 Enzymatic and Structural Roles: Intrinsic Excitability and Seizure Susceptibility

**Fig. S1.**
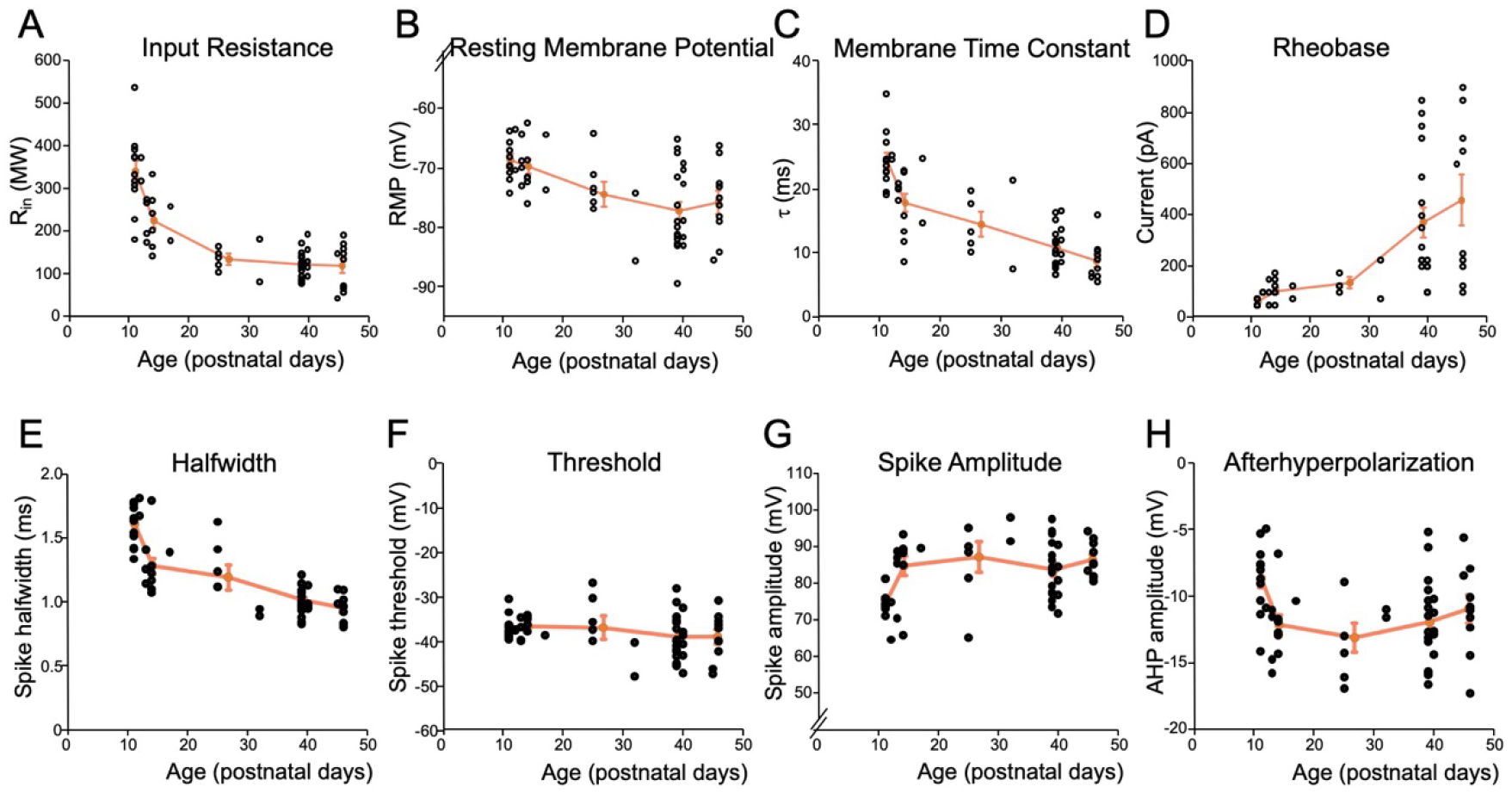
Properties of WT cells and their developmental time courses between P11 and P48. Combined data from WT cells of both strains showing input resistance (A), RMP (B), membrane time constant (C), rheobase (D), spike halfwidth (E), threshold (F), amplitude (G) and AHP (H). Black circles represent individual data points, orange circles and line are average values obtained by combining data from P11-12; P13-17; P28-32, P38-40 and >P45. There are developmental decreases in input resistance, RMP, membrane time constant and spike halfwidth; increases in rheobase and spike amplitude, and no consistent change in threshold and AHP.

**Fig. S2.**
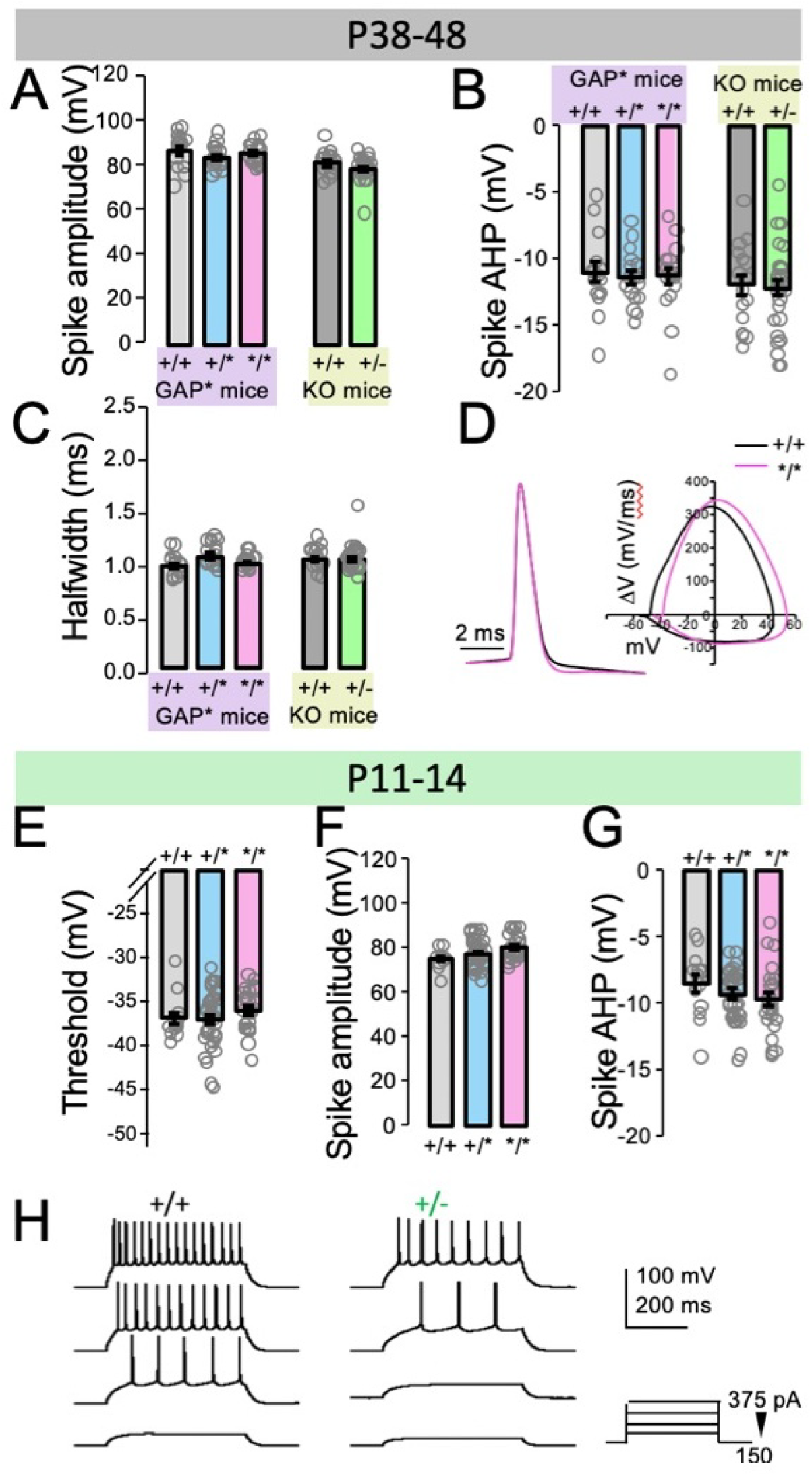
Intrinsic excitability in layer 2/3 pyramidal cells of GAP* and KO mice. A-C: Summary graphs showing pike amplitude (A), afterhyperpolarization (B) and halfwidth (C) in cells from GAP* and KO mice aged P38-48. D: Left: Representative action potentials from WT (black) and */* (magenta) mice, scaled to peak and aligned to threshold. Right: Phase plot (V vs ΔV) from the action potentials shown in the left. Halfwidth does not differ, but the */* cell has a higher spike threshold. E-F: Summary graphs showing pike threshold (E), amplitude (F) and afterhyperpolarization (G) in cells from GAP* mice aged P11-14. H: Example current clamp traces in from 500 ms current injections in +/+ (WT) and +/− (heterozygous knockout) cells, showing reduced spiking and increased rheobase in +/− cells. Current injections of +150, +225, +300 and +375 pA (bottom to top).

